# Automated data-intensive forecasting of plant phenology throughout the United States

**DOI:** 10.1101/634568

**Authors:** Shawn D. Taylor, Ethan P. White

**Affiliations:** School of Natural Resources and Environment, University of Florida Gainesville, FL, United States; Department of Wildlife Ecology and Conservation, University of Florida, Gainesville, FL, United States

**Keywords:** climate, budburst, flowering, phenophase, ecology, decision making

## Abstract

Phenology - the timing of cyclical and seasonal natural phenomena such as flowering and leaf out - is an integral part of ecological systems with impacts on human activities like environmental management, tourism, and agriculture. As a result, there are numerous potential applications for actionable predictions of when phenological events will occur. However, despite the availability of phenological data with large spatial, temporal, and taxonomic extents, and numerous phenology models, there has been no automated species-level forecasts of plant phenology. This is due in part to the challenges of building a system that integrates large volumes of climate observations and forecasts, uses that data to fit models and make predictions for large numbers of species, and consistently disseminates the results of these forecasts in interpretable ways. Here we describe a new near-term phenology forecasting system that makes predictions for the timing of budburst, flowers, ripe fruit, and fall colors for 78 species across the United States up to 6 months in advance and is updated every four days. We use the lessons learned in developing this system to provide guidance developing large-scale near-term ecological forecast systems more generally, to help advance the use of automated forecasting in ecology.

## Introduction

Plant phenology - the timing of cyclical and seasonal natural phenomena such as flowering and leaf out - influences many aspects of ecological systems (Chuine and Régnière 2017) from small scale community interactions (Ogilvie et al. 2017) to global scale climate feedbacks (Richardson et al. 2012). Because of the central importance of phenology, advanced forecasts for when phenological events will occur have numerous potential applications including: 1) research on the cascading effects of changing plant phenology on other organisms; 2) tourism planning related to flower blooms and autumn colors; 3) planning for sampling and application of management interventions by researchers and managers; and 4) agricultural decisions on timing for planting, harvesting, and application of pest prevention techniques. However, due to the challenges of automatically integrating, predicting, and disseminating large volumes of data, there are limited examples of applied phenology forecast systems.

Numerous phenology models have been developed to characterize the timing of major plant events and understand their drivers (Chuine et al. 2013). These models are based on the idea that plant phenology is primarily driven by weather, with seasonal temperatures being the primary driver at temperate latitudes (Basler 2016, Chuine and Régnière 2017). Because phenology is driven primarily by weather, it is possible to make predictions for the timing of phenology events based on forecasted weather conditions. The deployment of seasonal climate forecasts (Weisheimer and Palmer 2014), those beyond just a few weeks, provides the potential to forecast phenology months in advance. This time horizon is long enough to allow meaningful planning and action in response to these forecasts. With well established models, widely available data, and numerous use cases, plant phenology is well suited to serve as an exemplar for near-term ecological forecasting.

For decision making purposes, the most informative plant phenology forecasts will predict the response of large numbers of species and phenophases, over large spatial extents, and at fine spatial resolutions. The only regularly updated phenology forecast in current operation predicts only a single aggregated “spring index” that identifies when early-spring phenological events occur at the level of the entire ecosystem (not individual species) at a resolution of 1° lat/lon grid cells (Schwartz et al. 2013, Carrillo et al. 2018). Forecasting individual species and multiple phenological events at higher resolutions is challenging due to the advanced computational tools needed for building and maintaining data-intensive automatic forecasting systems (White et al. 2018, Welch et al. 2019). Automated forecasts requires building systems that acquire data, make model-based predictions for the future, and disseminate the forecasts to end-users, all in an automated pipeline (Dietze et al. 2018, White et al. 2018, Welch et al. 2019). This is challenging even for relatively small-scale single site projects with one to several species or response variables due to the need for advanced computational tools to support robust automation (White et al. 2018, Welch et al. 2019). Building an automated system to forecast phenology for numerous species at continental scales is even more challenging due to the large-scale data intensive nature of the analyses. Specifically, because phenology is sensitive to local climate conditions, phenology modeling and prediction should be done at high resolutions (Cook et al. 2010). This requires repeatedly conducting computationally intensive downscaling of seasonal climate forecasts and making large numbers of predictions. To make 4 km resolution spatially explicit forecasts for the 78 species in our study at continental scales requires over 90 million predictions for each updated forecast. To make the forecasts actionable these computational intensive steps need to be repeated in near real-time and disseminated in a way that allows end-users to understand the forecasts and their uncertainties (Dietze et al. 2018).

Here we describe an automated near-term phenology forecast system we developed to make continental scale forecasts for 78 different plant species. Starting December 1st, and updated every 4 days, this system uses the latest climate information to make forecasts for multiple phenophases and presents the resulting forecasts and their uncertainty on a dynamic website, https://phenology.naturecast.org/. Since the majority of plants complete budburst and/or flowering by the summer solstice in mid-June, this results in lead times of up to six months. We describe the key steps in the system construction, including: 1) fitting phenology models, 2) acquiring and downscaling climate data; 3) making predictions for phenological events; 4) disseminating those predictions; and 5) automating steps 2-4 to update forecasts at a sub-weekly frequency. We follow Welch et al. (2019)’s framework for describing operationalized dynamic management tools (ie. self-contained tools running automatically and regularly) and describe the major design decisions and lessons learned from implementing this system that will guide improvements to automated ecological forecasting systems. Due to the data-intensive nature of forecasting phenology at fine resolutions over large scales this system serves as a model for large-scale forecasting systems in ecology more broadly.

### Forecasting Pipeline

Welch et al. (2019) break down the process of developing tools for automated prediction into four stages: 1) Acquisition, obtaining and processing the regularly updated data needed for prediction; 2) Prediction, combining the data with models to estimate the outcome of interest; 3) Dissemination, the public presentation of the predictions; and 4) Automation, the tools and approaches used to automatically update the predictions using the newest data on a regular basis. We start by describing our approach to modeling phenology and then describe our approach to each of these stages.

### Phenology Modeling

Making large spatial scale phenology forecasts for a specific species requires species level observation data from as much of its respective range as possible (Taylor et al. 2019). We used data from the USA National Phenology Network (USA-NPN), which collects volunteer based data on phenological events and has amassed over 10 million observations representing over 1000 species. The USA-NPN protocol uses status-based monitoring, where observers answer ‘yes,’ ‘no,’ or ‘unsure’ when asked if an individual plant has a specific phenophase present (Denny et al. 2014). Phenophases refer to specific phases in the annual cycle of a plant, such as the presence of emerging leaves, flowers, fruit, or senescing leaves. We used the “Individual Phenometrics” data product, which provides pre-processed onset dates of individually monitored plants, for the phenophases budburst, flowering, and fall colors for all species with data between 2009 and 2017 (USA National Phenology Network 2018). We only kept “yes” observations where the individual plant also had a “no” observation within the prior 30 days and dropped any records where a single plant had conflicting records for phenotype status or more than one series of “yes” observations for a phenophase in a 12 month period. We built models for species and phenophase combinations with at least 30 observations (Figure 1, B) using daily mean temperature data at the location and time of each observation from the PRISM 4km dataset (PRISM Climate Group 2004). We also included contributed models of budburst, flowering, and/or fruiting for 5 species which were not well represented in the USA-NPN dataset (see Appendix S1: Table S2; Janet S. Prevéy, unpublished data, 2018, Prevéy et al. (In revision); Biederman et al. (2018)).

**Figure 1:**
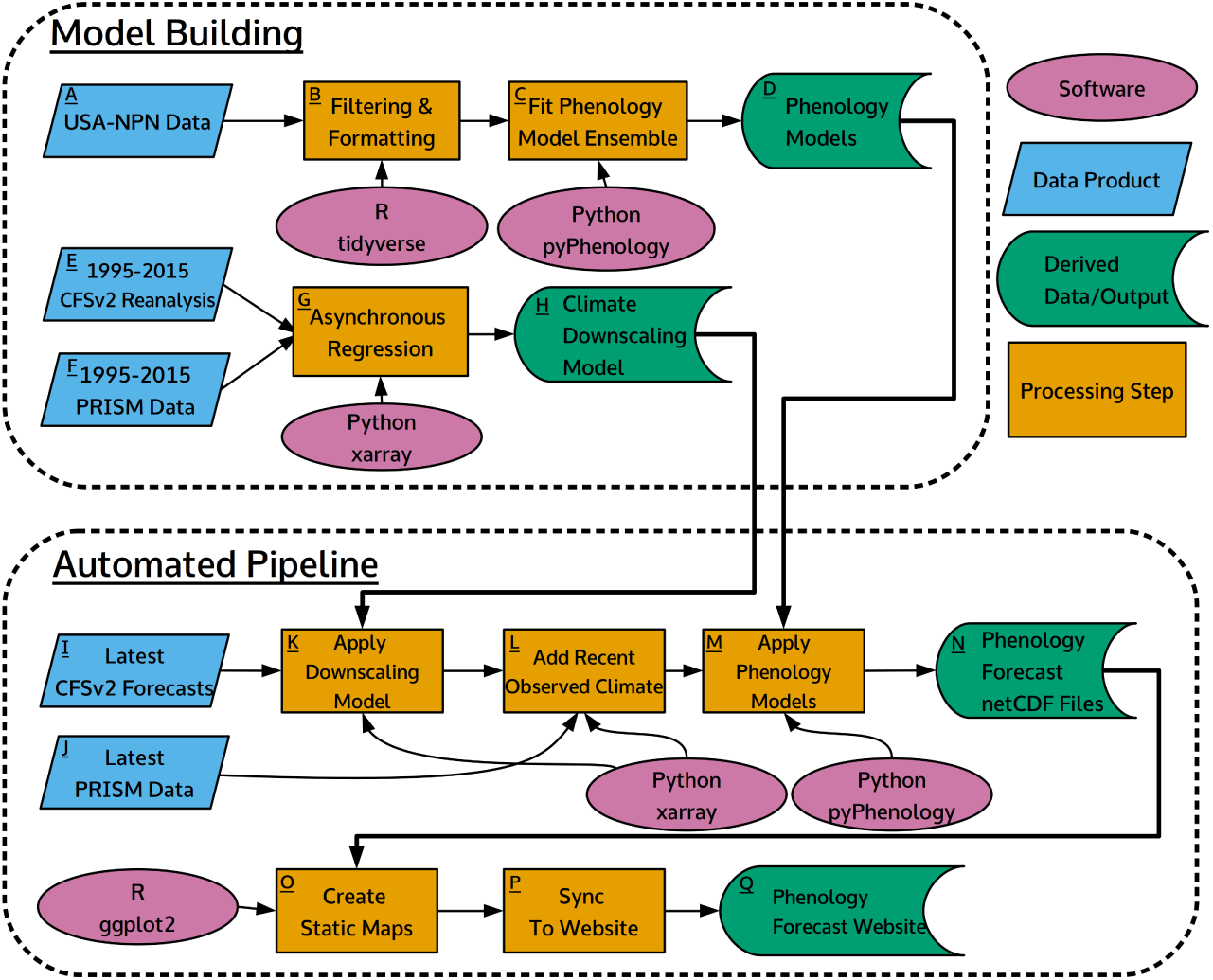
Flowchart of initial model building and automated pipeline steps. Letters indicate the associate steps discussed in the main text.

For each species and phenophase we fit an ensemble of four models using daily mean temperature as the sole driver (Figure 1, C). The general model form assumes a phenological event will occur once sufficient thermal forcing units accumulate from a specified start day (Chuine et al. 2013, Chuine and Régnière 2017). The specification of forcing units are model specific, but all are derived from the 24-hour daily mean temperature. In a basic model a forcing unit is the maximum of either 0 or the mean temperature above 0°C (ie. growing degree days). The amount of forcing units required, and the date from which they start accumulating are parameterized for each species and phenophase (see Appendix S1: Table S1). Ensembles of multiple models generally improve prediction over any single model by reducing bias and variance, and in a phenology context allow more accurate predictions to be made without knowing the specific physiological processes for each species (Basler 2016, Yun et al. 2017, Dormann et al. 2018). We used a weighted ensemble of four phenology models. We derived the weights for each model within the ensemble using stacking to minimize the root mean squared error on held out test data (100 fold cross-validation) as described in Dormann et al. (2018) (see Appendix S1: Sec. S1). After determining the weights we fit the core models a final time on the full dataset. Since individual process based phenology models are not probabilistic they do not allow the estimation of uncertainty in the forecasts. Therefore, we used the variance across the five climate models to represent uncertainty (see Prediction). Finally, we also fit a spatially corrected Long Term Average model for use in calculating anomalies (see Dissemination). This uses the past observations in a linear model with latitude as the sole predictor (see Appendix S1: Table S1).

In our pipeline 190 unique phenological models (one for each species and phenophase combination, see see Appendix S1: Table S2) needed to be individually parameterized, evaluated, and stored for future use. To consolidate all these requirements we built a dedicated software package written in Python, pyPhenology, to build, save, and load models, and also apply them to gridded climate datasets (Taylor 2018). The package also integrates the phenological model ensemble so that the four sub-models can be treated seamlessly as one in the pipeline. After parameterizing each model, its specifications are saved in a text based JSON file that is stored in a git repository along with a metadata file describing all models (Figure 1, D). This approach allows for the tracking and usage of hundreds of models, allowing models to be easily synchronized across systems, and tracking versions of models as they are updated (or even deleted).

### Acquisition and Downscaling of Climate Data

Since our phenology models are based on accumulated temperature forcing, making forecasts requires information on both observed temperatures (from Nov. 30 of the prior year up to the date a forecast is made) and forecast temperatures (from the forecast date onward). For observed data we used 4km 24-hour daily mean temperature from PRISM, a gridded climate dataset for the continental U.S.A. which interpolates on the ground measurements and is updated daily (PRISM Climate Group 2004). These observed data are saved in a netCDF file, which is appended with the most recent data every time the automated forecast is run. For climate forecasts we used the Climate Forecast System Version 2 (CFSv2; a coupled atmosphere-ocean-land global circulation model) 2-m temperature data, which has a 6-hour timestep and a spatial resolution of 0.25 degrees latitude/longitude (Saha et al. 2014). CFSv2 forecasts are projected out 9 months from the issue date and are updated every 6 hours. The five most recent climate forecasts are downloaded for each updated phenology forecast to accommodate uncertainty (see Prediction).

Because the gridded climate forecasts are issued at large spatial resolutions (0.25 degrees), this data requires downscaling to be used at ecologically relevant scales (Cook et al. 2010). A downscaling model relates observed values at the smaller scale to the larger scale values generated by the climate forecast during a past time period. We regressed these past conditions from a climate reanalysis of CFSv2 from 1995-2015 (Saha et al. 2010) against the 4km daily mean temperature from the PRISM dataset for the same time period (PRISM Climate Group 2004) to build a downscaling model using asynchronous regression (Figure 1, E-G). The CFSv2 data is first interpolated from the original 0.25 degree grid to a 4km grid using distance weighted sampling, then an asynchronous regression model is applied to each 4km pixel and calendar month (Stoner et al. 2013, see see Appendix S1: Sec. S2). The two parameters from the regression model for each 4 km cell are saved in a netCFD file by location and calendar month (Figure 1, H). This downscaling model, at the scale of the continental U.S.A., is used to downscale the most recent CFSv2 forecasts to a 4km resolution during the automated steps.

We used specialized Python packages to overcome the computational challenges inherent in the large CFSv2 climate dataset (Python Software Foundation 2003). The climate forecast data for each phenology forecast update is 10-40 gigabytes, depending on the time of year (time series are longer later in the year). While it is possible to obtain hardware capable of loading this dataset into memory, a more efficient approach is to perform the downscaling and phenology model operations iteratively by subsetting the climate dataset spatially and performing operations on one chunk at a time. We used the python package xarray (Hoyer and Hamman 2017), which allows these operations to be efficiently performed in parallel through tight integration with the dask package (Dask Development Team 2016). The combination of dask and xarray allows the analysis to be run on individual workstations, stand alone servers, and high performance computing systems, and to easily scale to more predictors and higher resolution data.

### Prediction

The five most recent downscaled climate forecasts are each combined with climate observations to make a five member ensemble of daily mean temperature across the continental USA (Figure 1, L). These are used to make predictions using the phenology model for each species and phenophase (Figure 1, M). Each climate ensemble member is a 3d matrix of latitude *×* longitude *×* time at daily timesteps extending from Nov. 1 of the prior year to 9 months past the issue date. The pyPhenology package uses this object to make predictions for every 4 km grid cell in the contiguous United States, producing a 2d matrix (latitude *×* longitude) where each cell represents the predicted Julian day of the phenological event. This results in approximately half a million predictions for each run of each phenology model and 90 million predictions per run of the forecasting pipeline. The output of each model is cropped to the range of the respective species (US Geological Survey 1999) and saved as a netCDF file (Figure 1, N) for use in dissemination and later evaluation.

An important aspect of making actionable forecasts is providing decision makers with information on the uncertainty of those predictions (Dietze et al. 2018). One major component of uncertainty that is often ignored in near-term ecological forecasting studies is the uncertainty in the forecasted drivers. We incorporate information on uncertainty in temperature, the only driver in our phenology models, using the CFSv2 climate ensemble (Figure 1, I; see Acquisition). The members of the climate ensemble each produce a different temperature forecast due to differences in initial conditions (Weisheimer and Palmer 2014). For each of the five climate members we make a prediction using the phenology ensemble, and the uncertainty is estimated as the variance of these predictions (see see Appendix S1: Sec. S1). This allows us to present the uncertainty associated with climate, along with a point estimate of the forecast, resulting in a range of dates over which a phenological event is likely to occur.

### Dissemination

To disseminate the forecasts we built a website that displays maps of the predictions for each unique species and phenophase (https://phenology.naturecast.org/; Figure 1 Q; Figure 2). We used the Django web framework and custom JavaScript to allow the user to select forecasts by species, phenophase, and issue date (Figure 2D). The main map shows the best estimate for when the phenological event will occur for the selected species (Figure 2A). Actionable forecasts also require an understanding of how much uncertainty is present in the prediction (Dietze et al. 2018), because knowing the expected date of an annual event such as flowering isn’t particularly useful if the confidence interval stretches over several months. Therefore we also display a map of uncertainty quantified as the 95% prediction interval, the range of days within which the phenology event is expected to fall 95% of the time (Figure 2C). Finally, to provide context to the current years predictions, we also map the predicted anomaly (Figure 2B). The anomaly is the difference between the predicted date and the long term, spatially corrected average date of the phenological event (Figure 1, O; see see Appendix S1: Table S1).

**Figure 2:**
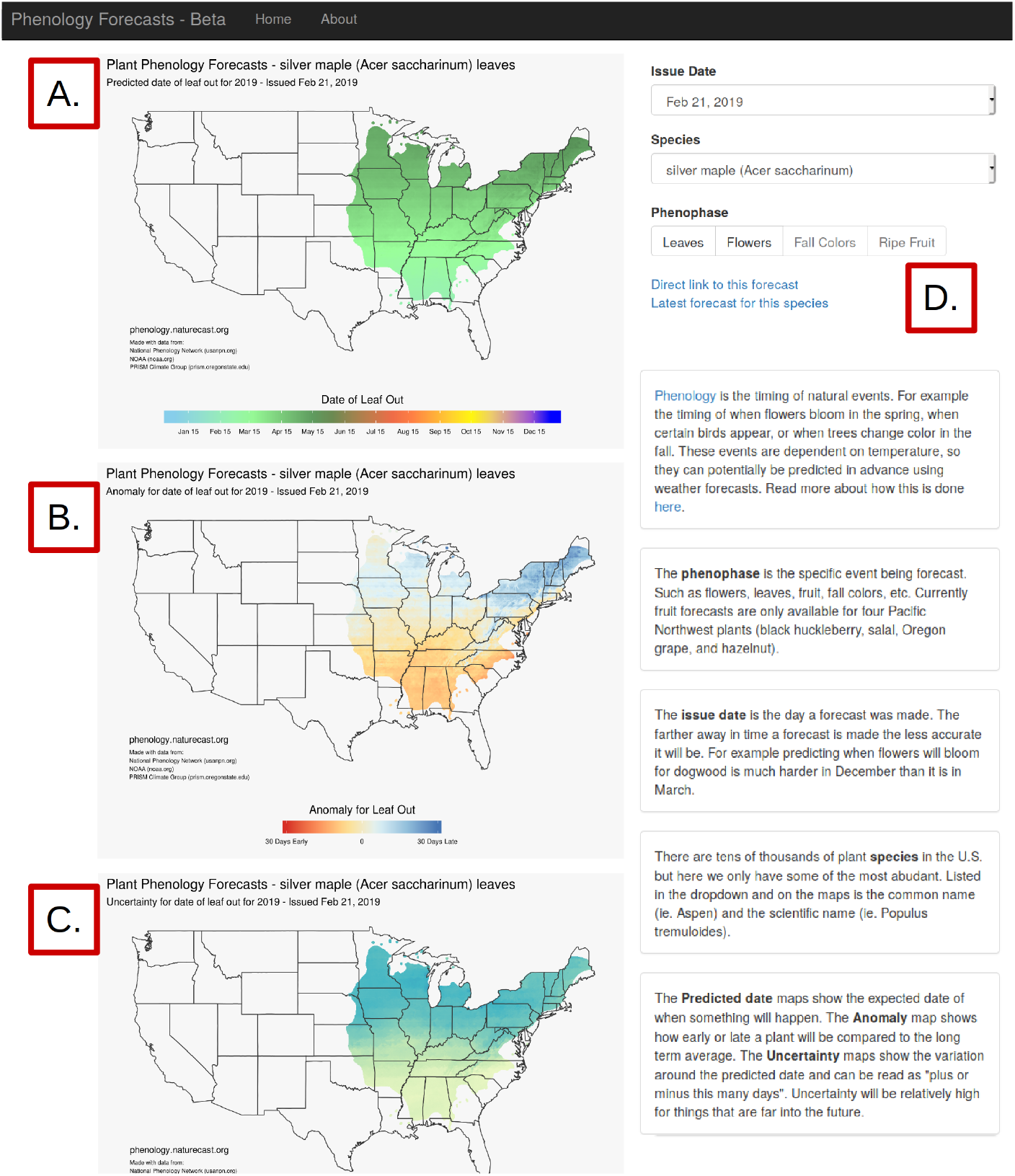
Screenshot of the forecast presentation website (http://phenology.naturecast.org) showing the forecast for the leaf out of *Acer saccharinum* in Spring, 2019, issued on Feburary 21, 2019. The maps represent the predicted date of leaf out (A), the anomaly compared to prior years (B), and the 95% confidence interval (C). In the upper right is the interface for selecting different species, phenophases, or forecast issue dates via drop down menus (D).

### Automation

All of the steps in this pipeline, other than phenology and downscaling model fitting, are automatically run every 4 days. To do this we use a cron job running on a local server. Cron jobs automatically rerun code on set intervals. The cron job initiates a python script which runs the major steps in the pipeline. First the latest CFSv2 climate forecasts are acquired, downscaled, and combined with the latest PRISM climate observations (Figure 1, I-L). This data is then combined with the phenology models using the pyPhenology package to make predictions for the timing of phenological events (Figure 1, M-N). These forecasts are then converted into maps and uploaded to the website (Figure 1, O-Q). To ensure that forecasts continue to run even when unexpected events occur it is necessary to develop pipelines that are robust to unexpected errors and missing data, and are also informative when failures inevitably do happen (Welch et al. 2019). We used status checks and logging to identify and fix problems and separated the website infrastructure from the rest of the pipeline. Data are checked during acquisition to determine if there are data problems and when possible alternate data is used to replace data with issues. For example, members of the CFSv2 ensemble sometimes have insufficient time series lengths. When this is the case that forecast is discarded and a preceding climate forecast obtained. With this setup occasional errors in upstream data can be ignored, and larger problems identified and corrected with minimal downtime. To prevent larger problems from preventing access to the most recent successful forecasts the website is only updated if all other steps run successfully. This ensures that user of the website can always access the latest forecasts.

Software packages used throughout the system include, for the R language, ggplot2 (Wickham 2016), raster (Hijmans 2017), prism (Hart and Bell 2015), sp (Pebesma and Bivand 2005), tidyr (Wickham and Henry 2018), lubridate (Grolemund and Wickham 2011), and ncdf4 (Pierce 2017). From the python language we also utilized xarray (Hoyer and Hamman 2017), dask, (Dask Development Team 2016), scipy (Jones et al. 2001), numpy (Oliphant 2006), pandas (McKinney 2010), and mpi4py (Dalcin et al. 2011). All code described is available on a GitHub repository (https://github.com/sdtaylor/phenology_forecasts). The code as well as 2019 forecasts and observations (see Evaluation) are also permanently archived on Zenodo (https://doi.org/10.5281/zenodo.2577452).

### Evaluation

A primary advantage of near-term forecasts is the ability to rapidly evaluate forecast proficiency, thereby shortening the model development cycle (Dietze et al. 2018). Phenological events happen throughout the growing season, providing a consistent stream of new observations to assess. We evaluated our forecasts (made from Dec. 1, 2018 thru May 1, 2019) using observations from the USA-NPN from Jan. 1, 2019 through May 8, 2019 and subset to species and phenophases represented in our system (Figure 3; USA National Phenology Network (2019)). This resulted in 1581 phenological events that our system had forecasts for (588 flowering events, 991 budburst events, and 2 fall coloring across 65 species, see see Appendix S1: Table S3). For each forecast issue date we calculated the root mean square error (RMSE) and average forecast uncertainty for all events and all prior issue dates. We also assessed the distribution of absolute errors 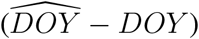 for a subset of issue dates (approximately two a month).

**Figure 3:**
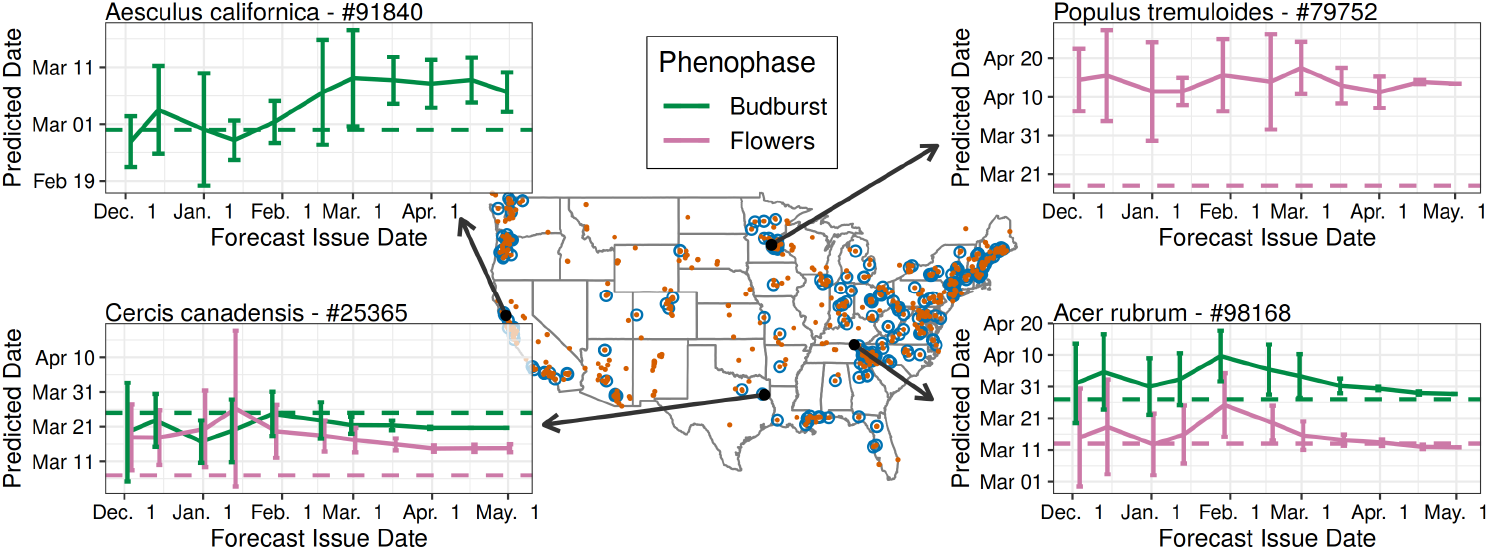
Locations of phenological events which have occurred between Jan. 1, 2019 and May 5, 2019 obtained from the USA National Phenology Network (blue circles), and all sampling locations in the same dataset (red points). Four individual plants are highlighted, with numbers indicating the USA National Phenology Network database ID. The solid line indicates the predicted event date as well as the 95% confidence interval for a specified forecast issue date, and the dashed line indicates the observed event date. The x-axis corresponds to the date a forecast was issued, while the y-axis is the date flowering or budburst was predicted to occur. For example: on Jan. 1, 2019 the *P. tremuloides* plant was forecast to flower sometime between March, 29 and April, 24 (solid lines). The actual flowering date was March 18 (dashed line).

Forecast RMSE and uncertainty both decreased for forecasts with shorter lead time (i.e. closer to the date the phenological event occurred), also known as the forecast horizon (Fig. 4; Petchey et al. (2015)). Forecasts issued at the start of the year (on Jan. 5, 2019) had a RMSE of 20.9 days, while the most recent forecasts (on May 5, 2019) had an RMSE of only 18.8 days. The average uncertainty for the forecasts were 7.6 and 0.2 days respectively for Jan. 5, and May 5. Errors were normally distributed with a small over-prediction bias (MAE values of 6.8 - 12.1, Fig. 5). This bias also decreased as spring progressed. These results indicate a generally well performing model, but also one with significant room for improvement that will be facilitated by the iterative nature of the forecasting system.

**Figure 4:**
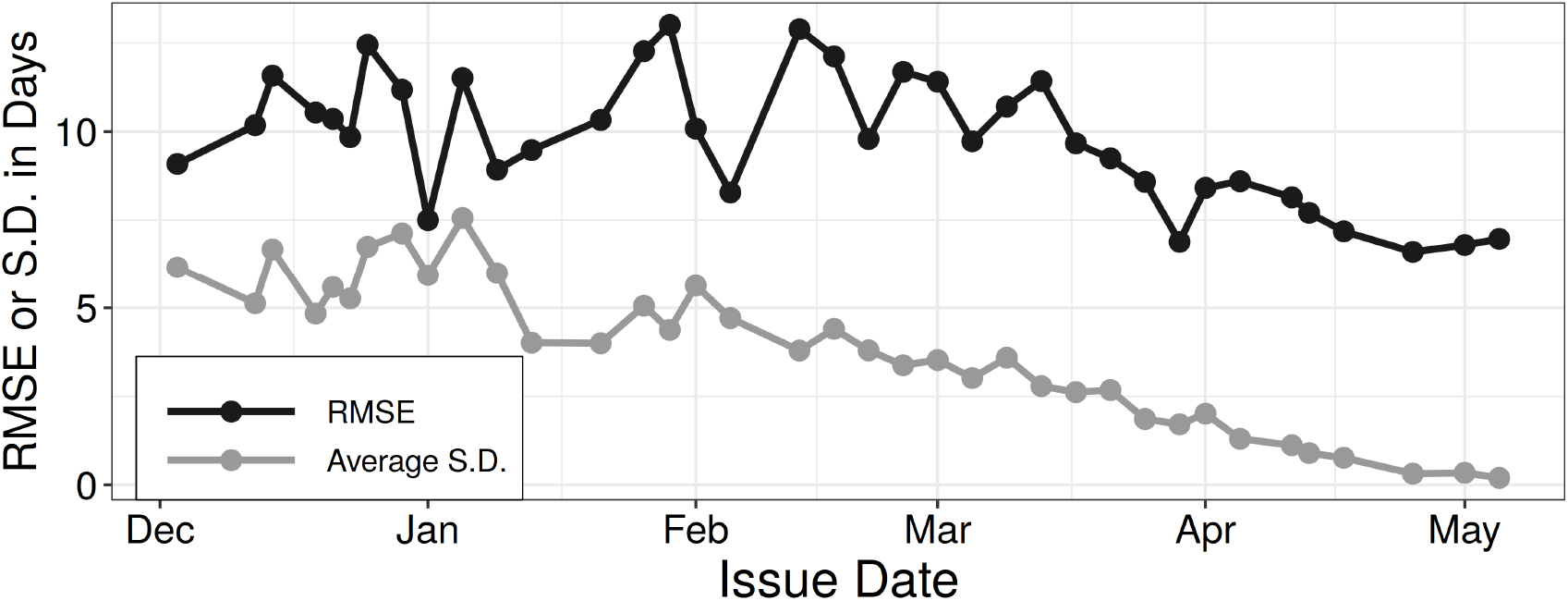
The root mean square error and the average uncertainty of forecasts issued between Dec. 2, 2018 and May 5, 2019 for 1581 phenological events representing 65 species.

**Figure 5:**
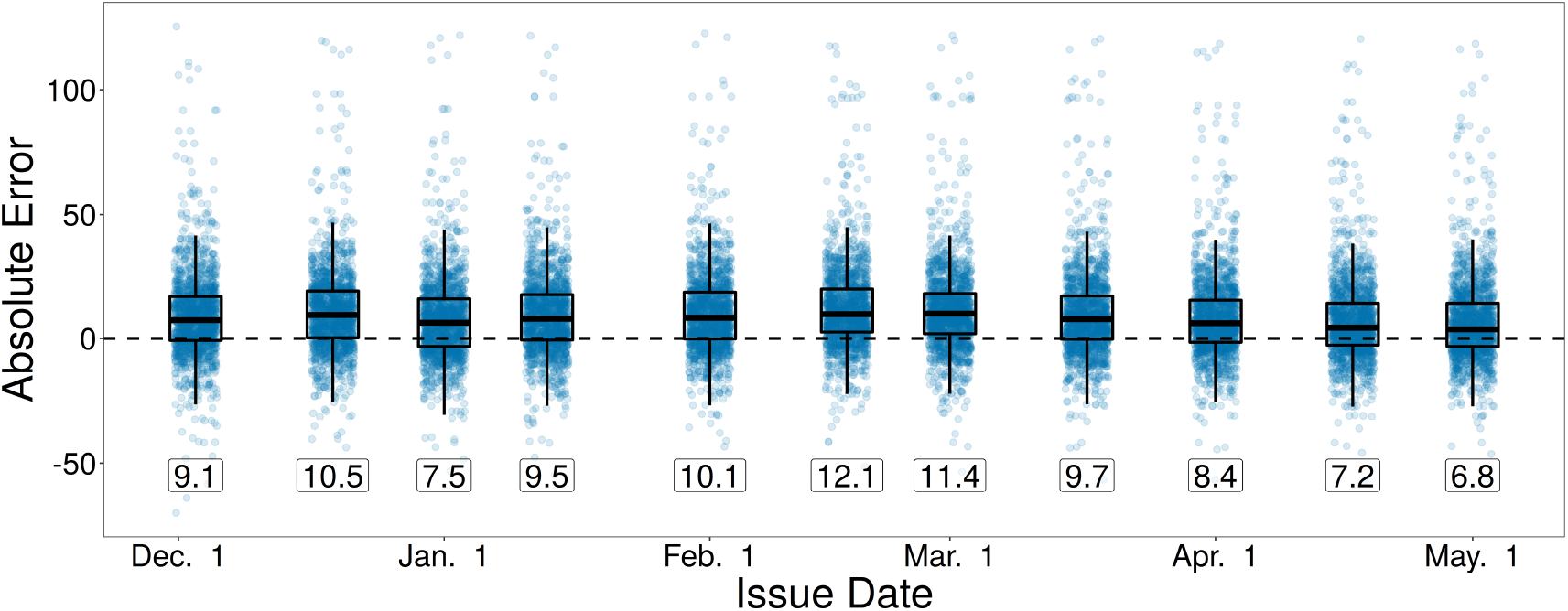
Distribution of absolute errors (prediction - observed) for 1581 phenological events for 11 selected issue dates. Labels indicate the mean absolute error (MAE).

## Discussion

We created an automated plant phenology forecasting system that makes forecasts for 78 species and 4 different phenophases across the entire contiguous United States. Forecasts are updated every four days with the most recent climate observations and forecasts, converted to static maps, and uploaded to a website for dissemination. We used only open source software and data formats, and free publicly available data. While a more comprehensive evaluation of forecast performance is outside the scope of this paper, we note that the majority of forecasts provide realistic phenology estimates across known latitudinal and elevational gradients (Figure 2), and forecast uncertainty and error decreases as spring progresses (Figure 4). While there is a bias from over-estimating phenological events, estimates were on-average within 2-3 weeks of the true dates throughout the spring season.

Developing automated forecasting systems in ecology is important both for providing decision makers with near real-time predictions and for improving our understanding of biological systems by allowing repeated tests of, and improvements to, ecological models (Dietze et al. 2018, White et al. 2018, Welch et al. 2019). To facilitate the development of ecological forecasts, we need both active development, descriptions, and discussion of a variety of forecasting systems. These discussions of the tools, philosophies, and challenges involved in forecast pipeline development will advance our understanding of how to most effectively build the systems, thereby lowering the entry barrier of operationalizing ecological models for decision making. Active development and discussion will also help us identify generalizable problems which can be solved with standardized methods, data formats, and software packages. Tools such as this can be used to more efficiently implement new ecological forecast systems, and facilitate synthetic analyses and comparisons across a variety of forecasts.

Automated forecasting systems typically involve multiple major steps in a combined pipeline. We found that breaking the pipeline into modular chunks made maintaining this large number of components more manageable (White et al. 2018, Welch et al. 2019). For generalizable pieces of the pipeline we found that turning them into software packages eased maintenance by decoupling dependencies and allowing independent testing. Packaging large components also makes it easier for others to use code developed for a forecasting system. The phenology modelling packge, pyPhenology, was developed for the current system, but is generalized for use in any phenological modelling study (Taylor 2018). We also found it useful to use different languages for different pieces of the pipeline. Our pipeline involved tasks ranging from automatically processing gigabytes of climate data to visualizing results to disseminating those results through a dynamic website. In such a pipeline no single language will fit all requirements, thus we made use of the strengths of two languages (Python and R) and their associate package ecosystems. Interoperability is facilitated by common data formats (csv and netCDF files), allowing scripts written in one language to communicate results to the next step in the pipeline written in another language.

This phenology forecasting system currently involves 190 different ensemble models, one for each species and phenological stage, each composed of 4 different phenology sub-models and their associated weights for a total of 760 different models. This necessitates having a system for storing and documenting models, and subsequently updating them with new data and/or methods over time. We stored the fitted models in JSON files (a open-standard text format). We used the version control system git to track changes to these text based model specifications. While git was originally designed tracking changes to code, it can also be leveraged for tracking data of many forms, including our model specifications (Ram 2013, Bryan 2018, Yenni et al. 2019). Managing many different models, including different versions of those models and their associate provenance, will likely be a common challenge for ecological forecasting (White et al. 2018) as one of the goals is iteratively improving the models.

The initial development of this system has highlighted several potential areas for improvement. First, the data-intensive nature of this forecasting system provides challenges and opportunities for disseminating results. Currently static maps show the forecast dates of phenological events across each species respective range. However this only answers one set of questions and makes it difficult for others to build on the forecasts. Additional user interface design, including interactive maps and the potential to view forecasts for a single location, would make it easier to ask other types of questions such as “Which species will be in bloom on this date in a particular location?”. User interface design is vital for successful dissemination, and tools such the python package Django used here, or the R packages Shiny and Rmarkdown provide flexible frameworks for implementation (White et al. 2018, Welch et al. 2019). In addition it would be useful to provide access to the raw data underlying each forecast. The sheer number of forecasts makes the bi-weekly forecast data relatively large, presenting some challenges for dissemination through traditional ecological archiving services like Dryad (https://datadryad.org) and Zenodo (https://zenodo.org). If stored as csv files every forecast would have generated 15 GB of data. We addressed this by storing the forecasts in compressed netCDF files, which are optimized for large-scale mutli-dimensional data and in our case are 300 times smaller than the csv files (50 MB/forecast).

In addition to areas for improvement in the forecasting system itself, its development has highlighted areas for potential improvement in phenology modeling. Other well-known phenological drivers could be incorporated into the models, such as precipitation and daylength. Precipitation forecasts are available from the CFSv2 dataset, though their accuracy is considerably lower than temperature forecasts (Saha et al. 2014). Other large-scale phenological datasets, such as remotely-sensed spring greenup could be used to constrain the species level forecasts made here (Melaas et al. 2016). Our system does not currently integrate observations about how phenology is progressing within a year to update the models. USA-NPN data are available in near real-time after they are submitted by volunteers, thus there is opportunity for data assimilation of phenology observations. Making new forecasts with the latest information not only on the current state of the climate, but also on the current state of the plants themselves would likely be very informative (Luo et al. 2011, Dietze 2017). For example, if a species is leafing out sooner than expected in one area it is likely that it will also leaf out sooner than expected in nearby regions. This type of data assimilation is important for making accurate forecasts in other disciplines including meteorology (Bauer et al. 2015, Carrassi et al. 2018). However, process based plant phenology models were not designed with data assimilation in mind (Chuine et al. 2013). Clark et al. (2014) built a bayesian hierarchical phenology model of budburst which incorporates the discrete observations of phenology data. This could serve as a starting point for a phenology forecasting model that incorporates data assimilation and allows species with relatively few observations to borrow strength from species with a large number of observations. The model from Clark et al. (2014) also incorporates all stages of the bud development process into a continuous latent state, thus there is also potential for forecasting the current phenological state of plants, instead of just the transition dates as is currently done in this forecast system.

Using recent advances in open source software and large-scale open data collection we have implemented an automated high resolution, continental scale, species-level phenology forecast system. Implementing a system of this scale was made possible by a new phenology data stream and new computational tools that facilitate large scale analysis with limited computing and human resources. Most recent research papers describing ecological forecast systems focus on only the modelling aspect (Chen et al. 2011, Carrillo et al. 2018, Van Doren and Horton 2018), and studies outlining implementation methods and best practices are lacking (but see White et al. 2018, Welch et al. 2019). Making a forecast system operational is key to producing applied tools, and requires a significant investment in time and other resources for data logistics and pipeline development. Major challenges here included the automated processing of large meteorological datasets, efficient application of hundreds of phenological models, and stable, consistently updated, and easy to understand dissemination of forecasts. By discussing how we addressed these challenges, and making our code publicly available, we hope to provide guidance for others developing ecological forecasting systems.

## Acknowledgments

This research was supported by the Gordon and Betty Moore Foundation’s Data-Driven Discovery Initiative through Grant GBMF4563 to E.P. White. We thank the USA National Phenology Network and the many participants who contribute to its Nature’s Notebook program.

