## Supplemental Information for "Automated data-intensive forecasting of plant phenology throughout the United States"

Phenology Model Descriptions

Description of the phenology model weighting

Description of the climate downscaling model

**Supplemental Tables 1-2**

### Phenology Model Descriptions

For all models, except the Linear and Naive models, the daily mean temperature $T_i$ is first transformed via the specified forcing equation. The cumulative sum of forcing is then calculated from a specific start date (either $DOY = 1$ or using the fitted parameter $t_1$, where $DOY$ is the day of year). The phenological event is estimated to be the $DOY$ where cumulative forcing is greater than or equal to the specified total required forcing (either $F^*$ or the specified equation). Parameters for each model are as follows: For the Linear model $\beta_1$ and $\beta_2$ are the intercept and slope, respectively and $T_{mean}$ is the average daily temperature between the spring start date and end using the parameters $Spring_{start}$ and $Spring_{start} + Spring_{length}$; in the Thermal Time model $F^*$ is the total accumulated forcing required, $t_1$ is the start date of forcing accumulation, and $T_{base}$ is the threshold daily mean temperature above which forcing accumulates; for the Alternating model $NCD$ is the number of chill days (daily mean temperature below 0°C) from $DOY = 1$ to the $DOY$ of the phenological event, $a$, $b$, and $c$ are the three fitted model coefficients; for the Uniforc model, is $F^*$ is the total accumulated forcing required, $t_1$ is the start date of forcing accumulation, and $b$ and $c$ are two additional fitted parameters which define the sigmoid function. For the Naive model $\beta_1$ and $\beta_2$ are the intercept and slope, respectively, for the average Julian day of a phenological event corrected for latitude. The Linear, Thermal Time, Alternating, andn Uniforc models are used in the primary forecast ensemble. The Long Term Average model does not use temperature data, and represents the long-term average, spatially correct average of a species/phenophase for use in annomoly calculations.

| Name | DOY Estimator | Forcing Equations | Reference |
|---|---|---|---|
| Linear | $DOY = \beta_1 + \beta_2 T_{mean}$ | - | - |
| Thermal Time | $\sum_{t=t_1}^{DOY} R_f(T_i) \geq F^*$ | $R_f(T_i) = max(T_i - T_{base}, 0)$ | (Réaumur 1735, Wang 1960, Hunter and Lechowicz 1992) |
| Alternating | $\sum_{t=1}^{DOY} R_f(T_i) \geq a + be^{cNCD(t)}$ | $R_f(T_i) = max(T_i - 5, 0)$ | (Cannell and Smith 1983) |
| Uniforc | $\sum_{t=t_1}^{DOY} R_f(T_i) \geq F^*$ | $R_f(T_i) = \frac{1}{1+e^{b(T_i - c)}}$ | (Chuine 2000) |

| Name | DOY Estimator | Forcing Equations | Reference |
|---|---|---|---|
| Long Term Average | $DOY = \beta_1 + \beta_2 Latitude$ | | |

### Description of the phenology model weighting

We used a weighted ensemble of the four models for each species and phenophase. The weights for each model within the ensemble were derived via stacking as described in Dormann et al. (2018). The steps for calculating weights are as followed:

1. Subset the phenology data into random training/testing sets.
2. Fit each core model on the training set.
3. Make predictions on the testing set.
4. Find the weights which minimize RMSE of the testing set.
5. Repeat 1-4 for 100 iterations.
6. Take the average weight for each model from all iterations as the final weight used in the ensemble. These will sum to 1.
7. Fit the core models a final time on the full dataset. Parameters derived from this final iterations will be used to make predictions, which are then used in the final weighted average.

The four phenology models are applied to each of the five members of the climate ensemble, with a final predicted value, $\widehat{DOY}_{forecast}$, derived as:

$$\widehat{DOY}_{forecast} = \frac{1}{5}\sum_{n=1}^{5}\sum_{i=1}^{4} w_i \widehat{DOY}_{n,i}$$

Where $n$ is the climate ensemble member, $i$ is the phenology model, $w$ is the phenology model weight, and $\widehat{DOY}$ the estimated Julian day.

Uncertainty is the 95% confidence interval of the estimates from the five climate ensembles:

$$2 * \sqrt{\frac{1}{5}\sum_{n=1}^{5}(\widehat{DOY}_n - \widehat{DOY}_{forecast})^2}$$

### Description of the climate downscaling model

The Climate Forecast System Version 2 (CFSv2) is a coupled atmosphere-ocean-land global circulation model maintained by the National Oceanic and Atmospheric Administration (NOAA)(Saha et al. 2014). The model tracks over 1000 global state variables of varying resolution and forecast length, such as ocean temperature and heights of pressure bands. Here we use the 2-meter temperature variable, which has a 6-hour timestep and a spatial resolution of 0.25 degrees latitude/longitude. The forecast is updated every 6 hours with the latest initial conditions and projected out 9 months.

The CFSv2 also has a reanalysis available. A climate reanalysis is a run of the full model over a prior time period with constant assimilation of known conditions. In practice this allows for analysis of state variables which are not able to be measured (such as the 500mb height over the arctic in winter). Here it allows us to build a downscaling model using the CFSv2 model's best estimate of past conditions of land surface temperature. These past conditions are regressed against finer grained "known" conditions from a different gridded dataset on a per pixel basis. We used the 2-m temperature output from the reanalysis from 1995-2015 as well as 4km daily mean temperature from the PRISM dataset (Climate 2004) to build a downscaling model using asynchronous regression (Figure 1, E-G). The model and theory are described in Stoner et al. (2013) and references therein. The CFSv2 data is first interpolated from the original 0.25 degree grid to a 4km grid using distance weighted sampling, then the following method is applied to each 4km pixel and calendar month.

1. Collect all daily mean temperature observations from 21 years of data from both the CFSv2 reanalysis and the PRISM dataset. This provides 588 - 641 points representing daily temperature for a single pixel and calendar month.
2. In addition to the data from each calendar month, also include data for the 14 days prior and 14 days following the calendar month, adding an addition 588 data points (21*(14+14)). This helps account for future novel conditions.

3. Order each dataset by their rank, such that the lowest value from the PRISM dataset is matched to the lowest value from the CFSv2 reanalysis.
4. Fit a linear regression model.

The two parameters from the regression model are saved in a netCFD file which can later be referenced by location and calendar month (Figure 1, H). This downscaling model, at the scale of the continental U.S.A., is used to downscale the most recent CFSv2 forecasts to a 4km resolution during the automated steps.

### Table S1

Species and their associated phenophases used in the forecast system. Note not all species have forecasts for all phenophases due to data availabilty. A * indicates a contributed model which was not built using USA-NPN data.

| | Species | Budburst | Fall Colors | Flowers | Ripe Fruits |
|---|---|---|---|---|---|
| 1 | Acacia greggii | | | ✓ | |
| 2 | Acer circinatum | ✓ | | ✓ | |
| 3 | Acer macrophyllum | ✓ | | ✓ | |
| 4 | Acer negundo | ✓ | ✓ | ✓ | |
| 5 | Acer pensylvanicum | ✓ | ✓ | ✓ | |
| 6 | Acer rubrum | ✓ | ✓ | ✓ | |
| 7 | Acer saccharinum | ✓ | ✓ | ✓ | |
| 8 | Acer saccharum | ✓ | ✓ | ✓ | |
| 9 | Aesculus californica | ✓ | ✓ | ✓ | |
| 10 | Alnus incana | ✓ | ✓ | ✓ | |
| 11 | Alnus rubra | ✓ | ✓ | | |
| 12 | Amelanchier alnifolia | ✓ | ✓ | ✓ | |
| 13 | Artemisia tridentata | | | ✓ | |
| 14 | Berberis aquifolium* | | | ✓ | ✓ |
| 15 | Betula alleghaniensis | ✓ | ✓ | ✓ | |
| 16 | Betula lenta | ✓ | ✓ | ✓ | |
| 17 | Betula nigra | ✓ | | ✓ | |
| 18 | Betula papyrifera | ✓ | ✓ | ✓ | |
| 19 | Carpinus caroliniana | ✓ | ✓ | ✓ | |
| 20 | Carya glabra | ✓ | ✓ | ✓ | |
| 21 | Celtis occidentalis | ✓ | | | |
| 22 | Cephalanthus occidentalis | ✓ | | ✓ | |
| 23 | Cercis canadensis | ✓ | ✓ | ✓ | |
| 24 | Chilopsis linearis | | | ✓ | |
| 25 | Clintonia borealis | | | ✓ | |
| 26 | Cornus florida | ✓ | ✓ | ✓ | |
| 27 | Cornus racemosa | ✓ | | | |
| 28 | Cornus sericea | ✓ | ✓ | ✓ | |
| 29 | Corylus cornuta* | | | ✓ | ✓ |
| 30 | Diospyros virginiana | ✓ | | | |
| 31 | Fagus grandifolia | ✓ | ✓ | ✓ | |
| 32 | Fouquieria splendens | | | ✓ | |
| 33 | Fraxinus americana | ✓ | ✓ | ✓ | |
| 34 | Fraxinus pennsylvanica | ✓ | ✓ | ✓ | |
| 35 | Gaultheria shallon* | | | ✓ | ✓ |
| 36 | Ginkgo biloba | ✓ | ✓ | ✓ | |
| 37 | Gleditsia triacanthos | ✓ | ✓ | ✓ | |
| 38 | Hamamelis virginiana | ✓ | ✓ | ✓ | |
| 39 | Ilex verticillata | ✓ | | ✓ | |
| 40 | Juglans nigra | ✓ | ✓ | ✓ | |
| 41 | Liquidambar styraciflua | ✓ | ✓ | ✓ | |
| 42 | Liriodendron tulipifera | ✓ | ✓ | ✓ | |
| 43 | Magnolia grandiflora | ✓ | | ✓ | |
| 44 | Maianthemum canadense | | | ✓ | |
| 45 | Nyssa sylvatica | ✓ | | ✓ | |
| 46 | Ostrya virginiana | ✓ | | ✓ | |
| 47 | Oxydendrum arboreum | ✓ | ✓ | ✓ | |
| 48 | Platanthera praeclara | ✓ | | ✓ | |
| 49 | Platanus racemosa | ✓ | | ✓ | |

| | Species | Budburst | Fall Colors | Flowers | Ripe Fruits |
|---|---|---|---|---|---|
| 50 | Populus deltoides | ✓ | ✓ | ✓ | |
| 51 | Populus fremontii | ✓ | ✓ | ✓ | |
| 52 | Populus tremuloides | ✓ | ✓ | ✓ | |
| 53 | Prunus americana | ✓ | | ✓ | |
| 54 | Prunus serotina | ✓ | ✓ | ✓ | |
| 55 | Prunus virginiana | ✓ | ✓ | ✓ | |
| 56 | Quercus agrifolia | ✓ | | ✓ | |
| 57 | Quercus alba | ✓ | ✓ | ✓ | |
| 58 | Quercus douglasii | ✓ | ✓ | ✓ | |
| 59 | Quercus gambelii | ✓ | ✓ | | |
| 60 | Quercus laurifolia | ✓ | | | |
| 61 | Quercus lobata | ✓ | ✓ | ✓ | |
| 62 | Quercus macrocarpa | ✓ | ✓ | ✓ | |
| 63 | Quercus palustris | ✓ | ✓ | ✓ | |
| 64 | Quercus rubra | ✓ | ✓ | ✓ | |
| 65 | Quercus velutina | ✓ | ✓ | ✓ | |
| 66 | Quercus virginiana | ✓ | | ✓ | |
| 67 | Rhododendron macrophyllum | ✓ | | ✓ | |
| 68 | Robinia pseudoacacia | ✓ | ✓ | ✓ | |
| 69 | Salix hookeriana | ✓ | ✓ | ✓ | |
| 70 | Salix lasiolepis | ✓ | | ✓ | |
| 71 | Sassafras albidum | ✓ | ✓ | ✓ | |
| 72 | Sorbus americana | ✓ | ✓ | ✓ | |
| 73 | Tilia americana | ✓ | ✓ | ✓ | |
| 74 | Ulmus americana | ✓ | ✓ | ✓ | |
| 75 | Umbellularia californica | ✓ | | ✓ | |
| 76 | Vaccinium corymbosum | ✓ | ✓ | ✓ | |
| 77 | Vaccinium membranaceum* | | | ✓ | ✓ |
| 78 | Yucca brevifolia | | | ✓ | |
| | **Total** | **67** | **47** | **72** | **4** |

### Table S2

Species and their associated phenophases evaluated from the 2019 season. Numbers indicate the total observations for the species and phenophase, with the mean Julian day in parentheses. Data are from the USA National Phenology Network from Jan. 1, 2019 - May 8, 2019.

| | Species | Budburst | Fall Colors | Flowers |
|---|---|---|---|---|
| 1 | Acer circinatum | 38 (93.1) | | 25 (112.4) |
| 2 | Acer macrophyllum | 10 (90.6) | | 6 (94.7) |
| 3 | Acer negundo | 14 (98.7) | | 13 (86.1) |
| 4 | Acer pensylvanicum | 6 (93.8) | | 2 (91) |
| 5 | Acer rubrum | 177 (93) | | 152 (86.1) |
| 6 | Acer saccharinum | 10 (107) | | 6 (99.3) |
| 7 | Acer saccharum | 56 (105.5) | | 30 (109.8) |
| 8 | Aesculus californica | 3 (46.3) | 2 (115) | 5 (118.2) |
| 9 | Alnus incana | 1 (107) | | |
| 10 | Alnus rubra | 6 (85.2) | | |
| 11 | Amelanchier alnifolia | 1 (83) | | |
| 12 | Betula alleghaniensis | 7 (114.3) | | 4 (122.5) |
| 13 | Betula lenta | 28 (99.2) | | 11 (106.5) |
| 14 | Betula nigra | 1 (104) | | 1 (87) |
| 15 | Betula papyrifera | 5 (112) | | 6 (113.8) |
| 16 | Carpinus caroliniana | 28 (82.1) | | 20 (80) |
| 17 | Carya glabra | 6 (77.2) | | 5 (112.8) |
| 18 | Celtis occidentalis | 4 (105.5) | | |
| 19 | Cephalanthus occidentalis | 9 (102) | | |
| 20 | Cercis canadensis | 22 (94.1) | | 24 (93) |
| 21 | Chilopsis linearis | | | 4 (119) |
| 22 | Cornus florida | 69 (89.7) | | 56 (101.8) |
| 23 | Cornus racemosa | 6 (102.5) | | |
| 24 | Cornus sericea | 9 (103.4) | | |
| 25 | Corylus cornuta | | | 4 (57.8) |
| 26 | Diospyros virginiana | 1 (124) | | |
| 27 | Fagus grandifolia | 45 (100.9) | | 10 (114.2) |
| 28 | Fouquieria splendens | | | 9 (97.7) |
| 29 | Fraxinus americana | 3 (109.7) | | |
| 30 | Fraxinus pennsylvanica | 2 (112.5) | | |
| 31 | Ginkgo biloba | 5 (108.6) | | 1 (111) |
| 32 | Hamamelis virginiana | 23 (104.3) | | |
| 33 | Ilex verticillata | 3 (115.3) | | |
| 34 | Juglans nigra | 5 (106.6) | | 4 (107.2) |
| 35 | Liquidambar styraciflua | 18 (75.2) | | 13 (89) |
| 36 | Liriodendron tulipifera | 41 (88.4) | | 14 (86.2) |
| 37 | Magnolia grandiflora | 5 (85.8) | | 7 (104.7) |
| 38 | Nyssa sylvatica | 17 (99) | | 1 (76) |
| 39 | Ostrya virginiana | 4 (91.5) | | |
| 40 | Oxydendrum arboreum | 8 (98.6) | | 1 (76) |
| 41 | Platanus racemosa | 5 (22) | | 3 (46) |
| 42 | Populus deltoides | 8 (111.8) | | 5 (110.6) |
| 43 | Populus fremontii | 2 (102) | | |
| 44 | Populus tremuloides | 10 (113) | | 11 (97.6) |

| | Species | Budburst | Fall Colors | Flowers |
|---|---|---|---|---|
| 45 | Prunus americana | 1 (111) | | 1 (112) |
| 46 | Prunus serotina | 30 (84.1) | | 10 (70.7) |
| 47 | Prunus virginiana | 6 (95.2) | | 1 (107) |
| 48 | Quercus agrifolia | 32 (68.7) | | 12 (83.2) |
| 49 | Quercus alba | 32 (100) | | 14 (112.7) |
| 50 | Quercus douglasii | 4 (81.8) | | 3 (108) |
| 51 | Quercus gambelii | 16 (114.2) | | |
| 52 | Quercus laurifolia | 9 (50.4) | | |
| 53 | Quercus lobata | 14 (63.9) | | 4 (81) |
| 54 | Quercus macrocarpa | 23 (108.5) | | 10 (120.5) |
| 55 | Quercus palustris | 4 (103) | | 2 (113) |
| 56 | Quercus rubra | 47 (105.8) | | 24 (112.9) |
| 57 | Quercus velutina | 4 (102.8) | | 3 (106) |
| 58 | Quercus virginiana | 10 (58) | | 11 (66.5) |
| 59 | Sassafras albidum | 6 (108) | | 8 (108) |
| 60 | Sorbus americana | 2 (124) | | |
| 61 | Tilia americana | 7 (101.9) | | |
| 62 | Ulmus americana | 8 (80.8) | | 2 (42) |
| 63 | Umbellularia californica | 5 (98.8) | | 5 (54.6) |
| 64 | Vaccinium corymbosum | 10 (81.1) | | 15 (90.4) |
| 65 | Yucca brevifolia | | | 10 (76.4) |
| | Total | 991 (93.8) | 2 (115) | 588 (94.8) |
